# Chronic Electrical Stimulation Promotes the Excitability and Plasticity of ESC-derived Neurons following Glutamate-induced Inhibition *In vitro*

**DOI:** 10.1101/354191

**Authors:** Charles-Francois V. Latchoumane, LaDonya Jackson, Mohammad .S Eslampanah Sendi, Kayvan F. Tehrani, Luke J. Mortensen, Steven L. Stice, Maysam Ghovanloo, Lohitash Karumbaiah

**Affiliations:** Regenerative Bioscience Center, ADS Complex, University of Georgia, Athens, Georgia; School of Chemical, Materials, and Biomedical Engineering, College of Engineering, University of Georgia, Athens, Georgia; School of Electrical and Computer Engineering, Georgia Institute of Technology, Atlanta, Georgia

**Keywords:** **Low Frequency Stimulation**, **Direct Current Stimulation**, **L-glutamate**, **Multi Electrode Arrays**, **Embryonic Stem cell-derived neurons**, **Traumatic brain injury**

## Abstract

Functional electrical stimulation (FES) is rapidly gaining traction as a therapeutic tool for mediating the repair and recovery of the injured central nervous system (CNS). However, the underlying mechanisms and impact of these stimulation paradigms at a molecular, cellular and network level remain largely unknown. In this study, we used embryonic stem cell (ESC)-derived neuron and glial cocultures to investigate network maturation following acute administration of L-glutamate, which is a known mediator of excitotoxicity following CNS injury. We then modulated network maturation using chronic low frequency stimulation (LFS) and direct current stimulation (DCS) protocols. We demonstrated that L-glutamate impaired the rate of maturation of ESC-derived neurons and glia immediately and over a week following acute treatment. The administration of chronic LFS and DCS protocols individually following L-glutamate infusion significantly promoted the excitability of neurons as well as network synchrony, while the combination of LFS/DCS did not. qRT-PCR analysis revealed that LFS and DCS alone significantly up-regulated the expression of excitability and plasticity-related transcripts encoding N-methyl-*D*-aspartate (NMDA) receptor subunit (NR2A), brain-derived neurotrophic factor (BDNF) and Ras-related protein (RAB3A). In contrast, the simultaneous administration of LFS/DCS down-regulated *BDNF* and *RAB3A* expression. Our results demonstrate that LFS and DCS stimulation can modulate network maturation excitability and synchrony following the acute administration of an inhibitory dose of L-glutamate, as well as an upregulation of NR2A, BDNF and RAB3A gene expression. Our study also provides a novel framework for investigating the effects of electrical stimulation on neuronal responses and network formation/repair after traumatic brain injury.

## INTRODUCTION

Direct current stimulation (DCS) has gained increasing attention in clinical and cognitive neuroscience applications due to it being a relatively inexpensive, non-invasive and easily parametrizable tool with minimal adverse effects ^1,2^. Transcranial DCS (tDCS) has been demonstrated to mediate therapeutic outcomes ^1,3,4^ in depression ^5–7^, dementia ^8,9^, pain ^10,11^ and central nervous system injury ^12–14^ In addition, tDCS applications extend to neuroprosthetics including improving working memory ^15^, motor and declarative learning ^16–19^ and decision making/addiction ^20–23^ In the healthy brain, the effects of tDCS and low frequency stimulation (LFS) on neuronal function and plasticity are thought to be mediated by changes in excitation/inhibition balance ^24–26^, modulation of the TrekB/BDNF pathway ^16^, NMDA receptor and intracellular calcium concentration ^27–29^ However, the underlying molecular, cellular and network mechanisms of action of tDCS remain largely unknown.

Traumatic brain injury (TBI) is a complex neuropathology commonly affecting healthy youth and war veterans ^30,31^, with a wide range of severity and phenotypic outcomes ^32^ The progression of TBI pathophysiology begins with the primary insult, which results in the acute impact induced mechanical damage to brain tissue. The secondary phase that follows involves inflammation, oxidative stress, metabolic dysregulation and excitotoxic release of neurotransmitters such as glutamate ^32–34^; resulting in significant brain tissue loss and functional impairments.. To this date, no therapeutic approaches have proven to be effective in treating TBI ^32^ tDCS has demonstrated significant promise in mediating functional recovery after TBI in preclinical studies ^35^, and in treating and rehabilitating brain function in TBI patients ^12,36,37^. The growing usage of tDCS as a therapeutic tool for TBI calls for a fundamental understanding of the underlying mechanisms of this new approach that would favor the development of more optimized protocols with temporally targeted applications ^38–40^.

Microelectrode array (MEA) recordings of neuronal cell cultures have been suggested as a promising proxy to the study of brain neural networks ^41^. MEA studies have helped characterize neuronal network maturation ^42^ and disruption under various pharmacological treatments ^43–45^, and could provide for realistic models to evaluate neuronal tissue development, disorders and injuries ^41^. The combination of DCS and MEA recordings to study the response of impaired neural networks remains underexplored, and could potentially help reveal important features and underlying mechanisms of action of electrical stimulation on brain tissue ^46^

Here, we used embryonic stem cell (ESC)-derived neuron and glia cocultures in combination with current stimulation protocols (i.e. DCS and LFS) and MEA recordings to investigate and modulate the response of neuronal networks to L-glutamate induced inhibition. Neurons and glia are both essential for the development and maintenance of synaptic connectivity within neural networks ^47,48^. Stem cells have begun to gain popularity as cells of choice to study neural network activity due to their ability to differentiate into both neurons and glia, and owing to their uniform clonal properties that can help minimize experimental variability ^49–51^. ESC-derived co-cultures containing astrocytes, oligodendrocytes, and a variety of neuron types can therefore serve as a model system to better understand baseline neural network function, and to investigate neuromodulatory mechanisms of recovering network activity after toxic exposure to excitatory neurotransmitters *in vitro* ^52–54^.

In this study, we present a model to study the impact of electric stimulation on the formation of ESC-derived neural networks following glutamate-induced inhibition. We quantified the impact of glutamate treatment on the rate of maturation in excitability and synchrony of neural networks as well as following three protocols based on chronic direct current stimulation: direct current stimulation (one time cathodal and daily anodal DCS; 6 days), low frequency stimulation (daily 0.1 Hz DCS stimulation; 5 days) and a combination of DCS and LFS. We quantified using qRT-PCR the change in excitability and synaptic function-related gene expression including *NR2A*, *NR2B*, *BDNF*, and *RAB3A*.

## MATERIALS AND METHODS

### 1. Cell culture

Mouse Hb9 ESC-derived neuron and glial cells (Aruna Biomedical, GA; Cat 7025) were cultured in AB2 basal neural medium (ArunA Biomedical, GA) and Dulbecco’s Modified Eagles Medium/F12 (DMEM/F12; Corning, NY) following a previously published protocol ^55^ Briefly, each well of 12 well 64 electrode per well containing MEA plates (Axion Biosystems, GA), or glass bottom petri dishes (Cellvis, CA) were coated with polyethylenimine (PEI) (Sigma-Aldrich, MO) and incubated for 1 h at 37°C and 5% CO_2_, then washed with deionized water and allowed to air dry overnight. The surfaces were subsequently coated with 20 μg/ml laminin (ThermoFisher Scientific, MA) for 2 h at 37°C and 5% CO_2_ immediately prior to seeding. The cells were subsequently seeded in triplicate on either glass bottom petri dishes, or in twelve well MEA plates. MEA plates (12-well, 1.43×1.43 mm surface) were seeded at a density of 40,000 cells per well, and glass-bottom petri dishes were seeded at a density of ~125,000 per well to maintain comparable cell seeding density to MEAs.

### 2. L-glutamate Inhibition Assays

Two weeks post-seeding, neuronal populations in glass-bottom petri dishes and MEA plates were treated with media only (controls) or with media containing 100 μM L-glutamate (Fisher Scientific, NH) and incubated at 37°C for a period of 20 min. Following this brief exposure, medium was removed and replaced with fresh complete growth medium devoid of excitotoxic agent and returned to the incubator. These cells were cultured for an additional week during which time electrophysiological assessments were made as described below. The cells were processed for immunocytochemical, and qRT-PCR analyses as described below at the experimental endpoint.

### 3. Electrophysiology Experiments and Analysis of Neural Metrics

Cellular electrophysiological activity was recorded for 5 min each at day 7, 14, and at day 21 from control and L-glutamate treated groups subjected to no stimulation, LFS-only, DCS-only, and LFS/DCS treatments. Recordings were acquired using the Maestro system (Axion Biosystems, GA) set at 37°C, and analyzed using Axis software (Axion Biosystems, GA). After data acquisition and recording, the media was removed and the wells were replenished with fresh media before returning the plate to the incubator.

Data obtained from each individual channel (electrode) was sampled with a gain setting of 1200x and a sampling rate of 12.5 kHz. The raw recorded local field potentials (LFP) were filtered using a Butterworth bandpass filter (200-3000 Hz cutoff frequencies) for all analyses. Event-triggered potentials were detected from the bandpass filtered data using a 6 × standard deviation root mean square (RMS) threshold on each channel and all triggered waveforms were saved on file along with raw local field potential data. An optional spike sorting was performed for electrode showing multi-unit activity using principal component analysis (PCA) decomposition of waveform and k-nearest neighbor (KNN) clustering using custom MATLAB scripts (Mathworks, Natick, MA). Offline analyses were performed using custom MATLAB scripts for spike properties (peak-to-valley amplitude and width, mean firing rate, mean bursting rate) and network properties (# of units per well, # of active electrode per well, # of bursting cells per well, mean population firing rate, mean population bursting rate, event synchronization and spike train crosscorrelation peak).

### 4. Measures of excitability

For each recording electrode (64 channels per wells), we estimated the number of triggered spikes and classified the electrode as active if at least 5 spikes/min were observed over the 5 minutes of recording ^44^ The total number of active electrodes was then derived for each wells.

The weighted mean firing rate (wMFR) can estimate the overall population excitability and connectivity, and was derived as the mean firing rate of a well, averaged over the active electrodes (wMFR is express as spikes per sec per active electrode). The instantaneous wMFR is derived after binning the activity of all electrodes (bin size: 100 msec) and is estimated as spike per bin and normalized to the number of active electrodes (Figure 3A).

For each spiking units/electrode, bursting was estimated following low-threshold rebound burst criteria: 50 msec of no spiking (silencing period), followed by at least 3 spikes with an inter-spike interval lower than 5 msec; the maximum inter-spike interval was set to 20 msec within one burst. The number of bursting cells per well was estimated from the number of spiking units/electrodes showing low-threshold burst.

### 5. Measures of Synchrony

Population burst estimation was based on the instantaneous wMFR and the instantaneous count of active electrode per time bin (CellCount per Bin) ^56–58^ A population burst was counted if: 1) at least 30% of active electrodes spiking together within a 100 msec bin, and 2) the wMFR for that same bin was higher than 0.5 spikes/bin/electrodes. The start and stop of the burst were estimated from a threshold crossing. The threshold value was for burst start and stop was estimated from the cumulative histogram of the product of instantaneous wMFR and CellCountBin (50%). The mean population burst firing rate (mBFR) was then estimated as the count of burst event over the entire recording duration (burst event/sec).

Event synchronization ^59,60^ was obtained from the average event synchronization value over all unique pairs of spiking units/electrodes (only active electrodes were considered for the estimation). Briefly, for a single pair of electrodes, the event synchronization was derived from the number of spiking events that appeared within a short window of time (quasi-simultaneous) and normalized to be: a) symmetrical (i.e. no directionality), and b) unitary for identical spike trains (i.e. spikes showing identical and asynchronous spike train would have an event synchronization value of 1 and 0, respectively).

The cross-correlation based synchrony (cross-correlation peak) is derived from the average cross-correlation peak over all unique pairs of active electrodes ^61,62^. For each unique pair, the cross-correlogram histogram (range [−10, 10] sec; bin size = 100 msec) is estimated and normalized (normalization based on the square root of the product of the two spike trains length). The cross-correlation peak is the value of the normalized cross-correlogram at lag = 0.

### 6. Low Frequency and Direct Current Stimulation

LFS was administered through the electrodes in each well using the Maestro system. The Axis software was programmed to deliver monophasic anodal LFS at 10 μA and at a frequency of 0.1 Hz for 15 minutes every day from days 15-19 (Figure 6A) ^16^. Monophasic cathodal and anodal direct current stimulation (DCS) was administered from a battery powered external device that contained variable resistors set to deliver 10 μA through a Keithley Series 2280 power supply (Keithley Instruments, OH) and a floating ground (Supplementary Figure 3). Cathodal DCS was administered at the onset of excitotoxicity on Day 14 (15min), and anodal DCS was administered from day 15-19 for 15min/day (Figure 6A). The initial Cathodal DCS stimulation has been suggested to reduce the extent of glutamate-induced hyper-excitability and cytotoxicity, and its impact on brain recovery following injury ^63,64^. Day 15 to 19 stimulations (LFS and DCS anodal) were designed to entrain the neural network into an excitable state, hypothetically favorable for its maturation ^65,66^.

### 7. Immunocytochemistry

Cells seeded in glass bottom petri-dishes were fixed in 4% paraformaldehyde containing 0.4 M sucrose in phosphate buffered saline (PBS) for 20 minutes at 1 and 3 weeks post-seeding. They were then washed thrice in PBS and permeabilized in blocking buffer (PBS containing 4% goat serum, 0.5% Triton-×100) for 1 h. Cells were then incubated with blocking buffer containing primary antibodies overnight at 4°C. Primary antibodies specific to β-III tubulin (1:200; Millipore, MA), Sox-1 (1:500; R&D Systems, MN), GFAP (1:500; Dako Agilent, CA) and O4 (1:500; R&D systems, MN) were used to mark neurons, neural stem cells, astrocytes, and oligodendrocytes, respectively. The next day, the cells were rinsed thrice with PBS, and incubated with blocking buffer for 1 h. Appropriately matched 555 anti-Mouse IgM (1:220; ThermoFisher Scientific, MA), 647 Goat anti-Mouse IgG1 (1:220; ThermoFisher Scientific, MA), 555 Goat antiRabbit IgG (1:220; ThermoFisher Scientific, MA), and 488 anti-Chicken IgY (1:220; ThermoFisher Scientific, MA) secondary antibodies in blocking buffer were then applied for 1 h. The cells were then again washed thrice with PBS then incubated in a nuclear stain (NucBlue; ThermoFisher Scientific, MA) for 5 min, and washed thrice with PBS. The cells were mounted with fluoromount-G (SouthernBiotech, AL), coverslipped, and imaged using a Leica DMIRBE fluorescence microscope (Leica Microsystems Inc, IL).

### 8. qRT-PCR Analysis

Total RNA was isolated using RNeasy Plus Mini kit (Qiagen, CA) on day-21 from LFS or DCS stimulated and unstimulated cells belonging to control (no glutamate exposure) and treated (exposed to 100 μM L-glutamate), respectively (Table 2). Total RNA was quantified using a NanoDrop 8000 (ThermoFisher Scientific, MA), and cDNA was synthesized using the RT First Strand cDNA synthesis kit (Qiagen, CA). A total of 100 ng total RNA equivalent of cDNA template was used in 25 μL qRT-PCR reactions for each treatment group along with SYBR green dye (Qiagen, CA), and pre-validated primers targeting mouse *NR2A*, *NR2B*, *BDNF*, *RAB3A*; and the endogenous housekeeping genes *GAPDH* and *HPRT1* (Qiagen, CA) and amplified using a ABI 7900HT qRT-PCR instrument (Applied Biosystems, CA) using conditions described previously ^67,68^. Each sample was assayed in triplicate using cycle conditions: 95°C for 10 minutes, 40 cycles of 95 °C for 15 seconds, and 60 °C for 1 minute, followed by a melting curve analysis. Relative quantitative gene expression was determined using the ΔΔCT method. The fold increase or decrease in target gene expression was calculated after normalization to media-only control and against endogenous controls for each sample and then presented as relative units.

### 9. Statistical Analysis

Analyses of immunocytochemical staining and cell quantification was performed using Volocity software (PerkinElmer, MA). Unless stated otherwise all results are expressed as mean +/− SEM. Repeated measure ANOVA, two-way and one-way ANOVA were performed for longitudinal and group difference test with post-hoc correction for multiple comparisons when appropriate (Holm-Sidak). When necessary (normality test fail or equality of variance test fail), nonparametric alternative test were performed, i.e. Kruskal-wallis test (with post-hoc Dunn-Sidak correction), Wilcoxson ranksum test and Signrank test. Distribution comparisons were performed either using the Kolmogorov-Smirnov test (Lillie’s correction), Shapiro-Wilk test, or the Chi-square test goodness-of-fit test. Differences were considered statistically significant if p < 0.05. All statistical tests were performed using either SigmaPlot (Systat Software inc., CA) or MATLAB^®^ statistical tool box (MathWorks Inc., MA).

## RESULTS

### ESC-derived neurons and glia mimic the cellular composition of mature neural tissue

In this study, we used mouse ESC-derived neurons and glia in order to investigate the maturation and composition of neural tissue (See Materials and Method section for details).

We found a characteristic differentiation (Figure 1A) of the ESC-derived cells as soon as week 1 into β-III tub+ (neuronal specific marker) neurons and SOX-1+ (a neuronal marker for cell with progenitor stem cell origin) cells. The ESC-derived cells also showed positive differentiation into oligodendrocytes (Figure 1-B, O4+) and astrocytes (Figure 1-B, GFAP+). These observations were confirmed by cell density quantification (Supp. Figure 1). We found that over three weeks of maturation, ESC-derived neurons had an increased differentiation toward βIII-tub+ neurons that plateaued at week 2 (Supp. Figure 1-A) whereas ESC-derived glia had a steady differentiation toward O4+ and GFAP+ cells up until week3 (Supp. Figure 1-B).

The results indicate that the ESC-derived neurons and glia mature to a dynamically increasing density of βIII-tub+ neurons, O4+ oligodendrocytes, and GFAP+ astrocytes over a period of three weeks *in vitro*.

**Figure 1:**
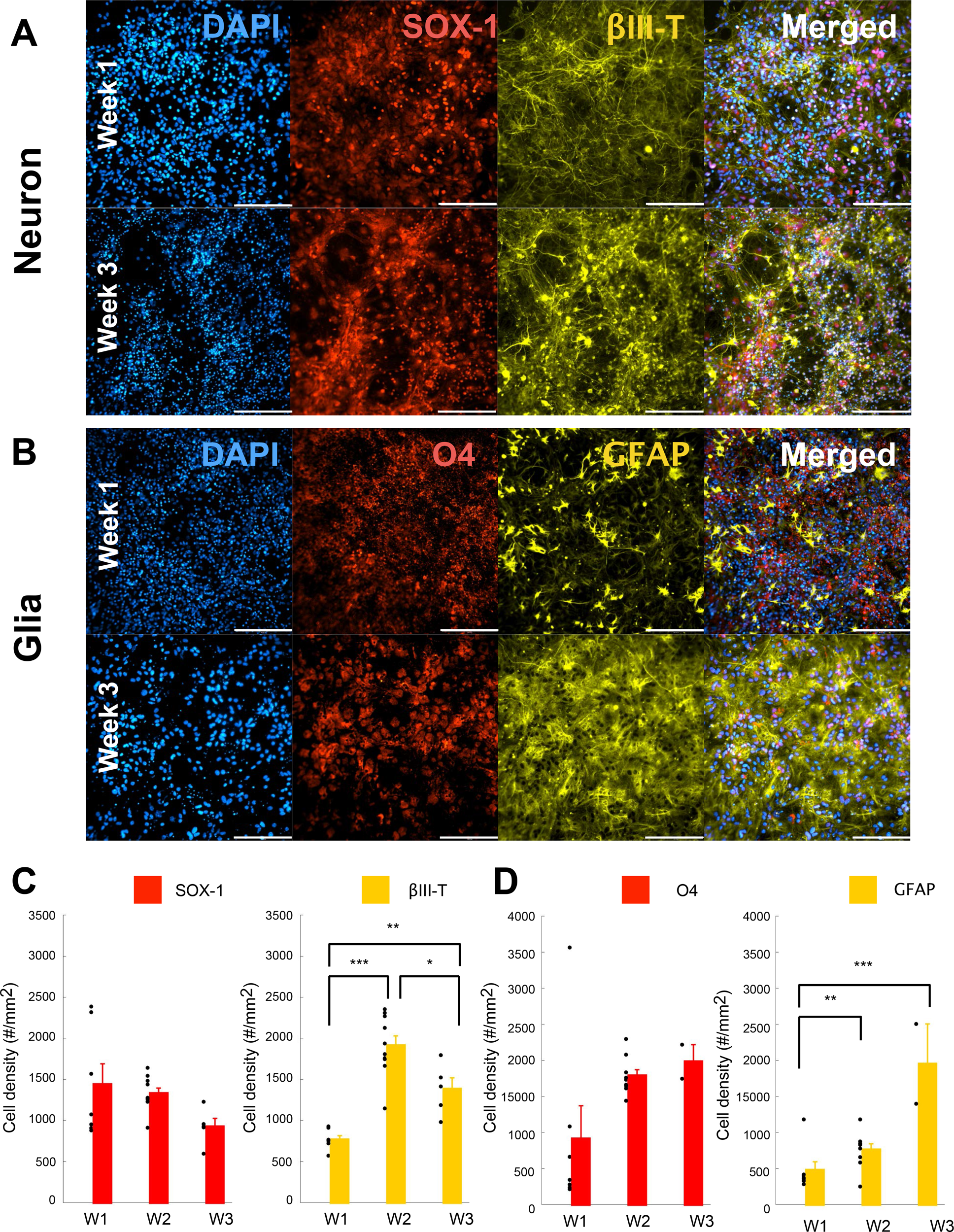
ESC-derived neurons and glia matured into a neural tissue. (**A**) ESC-derived neurons population for the control condition (CTR, no treatment) at week 1 (top panels) and week 3 (bottom panels). From left to right panels are shown DAPI, SOX-1 (Neural progenitor origin marker), βIII-Tubulin (Neuronal marker) and merged image for all 3 fluorescent markers. The scale bar is for 200 μm. (**B**) ESC-derived glial population for the control condition (CTR, no treatment) at week 1 (top panels) and week 3(bottom panels). From left to right panels are shown DAPI, O4 (Oligodendrocyte marker), GFAP (Glial marker) and merged image for all 3 fluorescent markers. The scale bar is for 200 μm. (**C**) Neuronal cell density (ȣ/mm2) estimated for the control condition (CTR) over week1 through week3. Quantification for the SOX-1 marker (Neurogenesis marker; left panel) and Beta-III tubulin (Neuronal marker; right panel). Data is shown as scatter plot of individual plate quantification and bar plot using mean and s.e.m. * is for p<0.05, ** is for p<0.01 and *** is for p<0.001 using post-hoc multiple comparison with Holm-Sidak correction. (**D**) Glia cell density (#/mm2) estimated for the control condition (CTR) over week1 through week3. Quantification for the GFP marker (Differentiated motor neurons; left panel), O4 marker (Oligodendrocyte marker; middle panel) and GFAP (Glial marker; right panel). Data is shown as scatter plot of individual plate quantification and bar plot using mean and s.e.m. * is for p<0.05, ** is for p<0.01 and *** is for p<0.001 using post-hoc multiple comparison with Holm-Sidak correction.

### ESC-derived neurons and glia form interconnected neural networks

In order to characterize the integrity and functionality of the neural network formed using the ESC-derived neurons and glia, we used MEA recording assays (Figure 2-B). ESC-derived cells were seeded at day 1 in each well of 12-well MEA plates, and recordings were acquired at weeks 1 (7 days following seeding), 2 and 3 (Figure 2-A).

We observed that the distribution of spike width (peak-to-valley width: 553±150 μsec) and spike amplitude (peak-to-valley amplitude: 37±12 μVolt) was normal (Figure 2C) and typical of neuronal spike waveforms (Figure 2D) ^69,70^.

We quantified the functional connection between active electrodes/units using event synchronization ^59,60^ and cross-correlation peak measures ^61,62^. As expected, ESC-derived neurons and glia formed networks that exhibited synchronous activity (Figure 2E), and with a high tendency for network bursting (See materials and method for detail).

Consistently with the observed neuronal differentiation, the activity observed at each electrode increased over time (Supplementary Figure 1). The number of active electrodes (i.e. electrode with spiking rate > 5 spikes/min) per MEA (1 well, 64 channels) showed a significant increase from week 1 through week 3 (Figure 3; repeated-measure ANOVA, time: p < 0.001; post-hoc Holm-Sidak correction: pw2-w3 < 0.01 and pw1-w3 < 0.001). ESC-derived neural networks showed increasing synchrony over time (Figure 3BC; repeated measure ANOVA, time: p < 0.001 for both event synchronization and cross-correlation peak). In addition, we observed that over time, the network bursting rate showed a marginal increase (repeated measure ANOVA, time: p = 0.07) and the mean burst duration significantly decreased over time (repeated measure ANOVA, time: p < 0.001).

Altogether, these results indicate that the ESC-derived neuronal and glial cells mature into a connected neural network that demonstrate a temporal increase in synchronization and population bursting.

**Figure 2:**
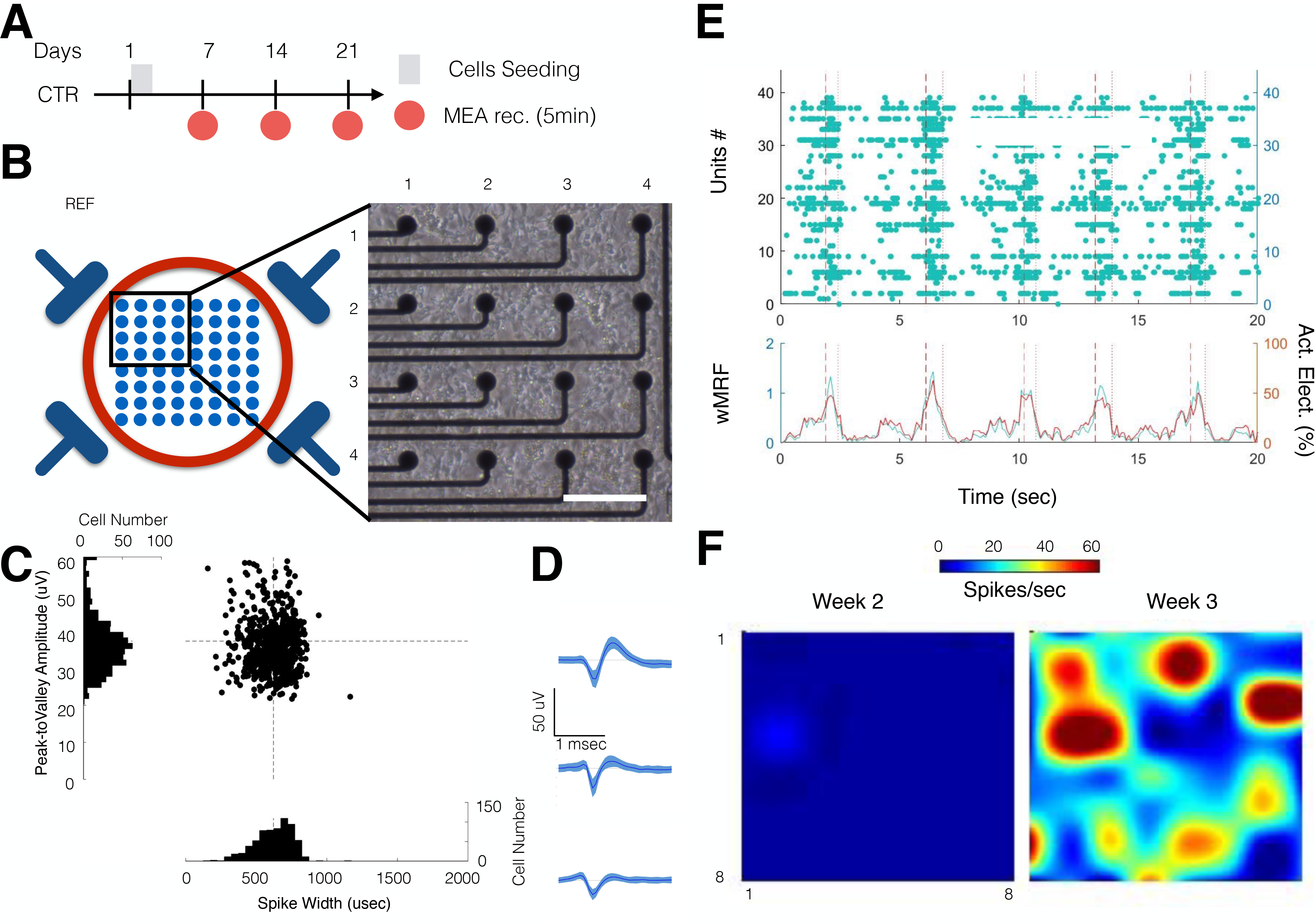
ESC-derived neurons and glia form functional neural networks on MEAs. (**A**) Experimental schedule for neural stem cell seeding and culture. For the control group, recording were performed from week1 to week 3. (**B**) Representative micro-electrode array (MEA) recording setup. Schematic of the bottom of the well with 4 reference electrodes and 8×8 electrode array (left). Top left quadrant of a 64-channel MEA plate, 2 weeks after stem cells seeding (right; bar is for 100um). (**C**) Scatter distribution of the spike width (μsec; x-axis) against the peak-to-valley amplitude (μVolt; y-axis) for individual unit recorded for the CTR at week 3. The distribution (cell count per bin) is displayed as a projection of each axis. The dashed black line represents the average of spike width (vertical) and peak-to-valley amplitude (horizontal). (**D**) Representative average spike wave forms obtain from 3 electrodes at week 3 for the control group. Data is shown as mean and s.e.m. (**E**) Raster plot of one well recorded at week 3 (top panel). The population instantaneous firing rate (left y-axis in green; wMFR, spikes/bin/electrode) and the percentage of active electrode per bin (right y-axis, in red; active electrode in %). The vertical dashed and dotted lines indicate the start and stop of a detected population burst, respectively. (**F**) Heatmap showing the evolution of the average activity in a control well from week 2 to week 3.

**Figure 3:**
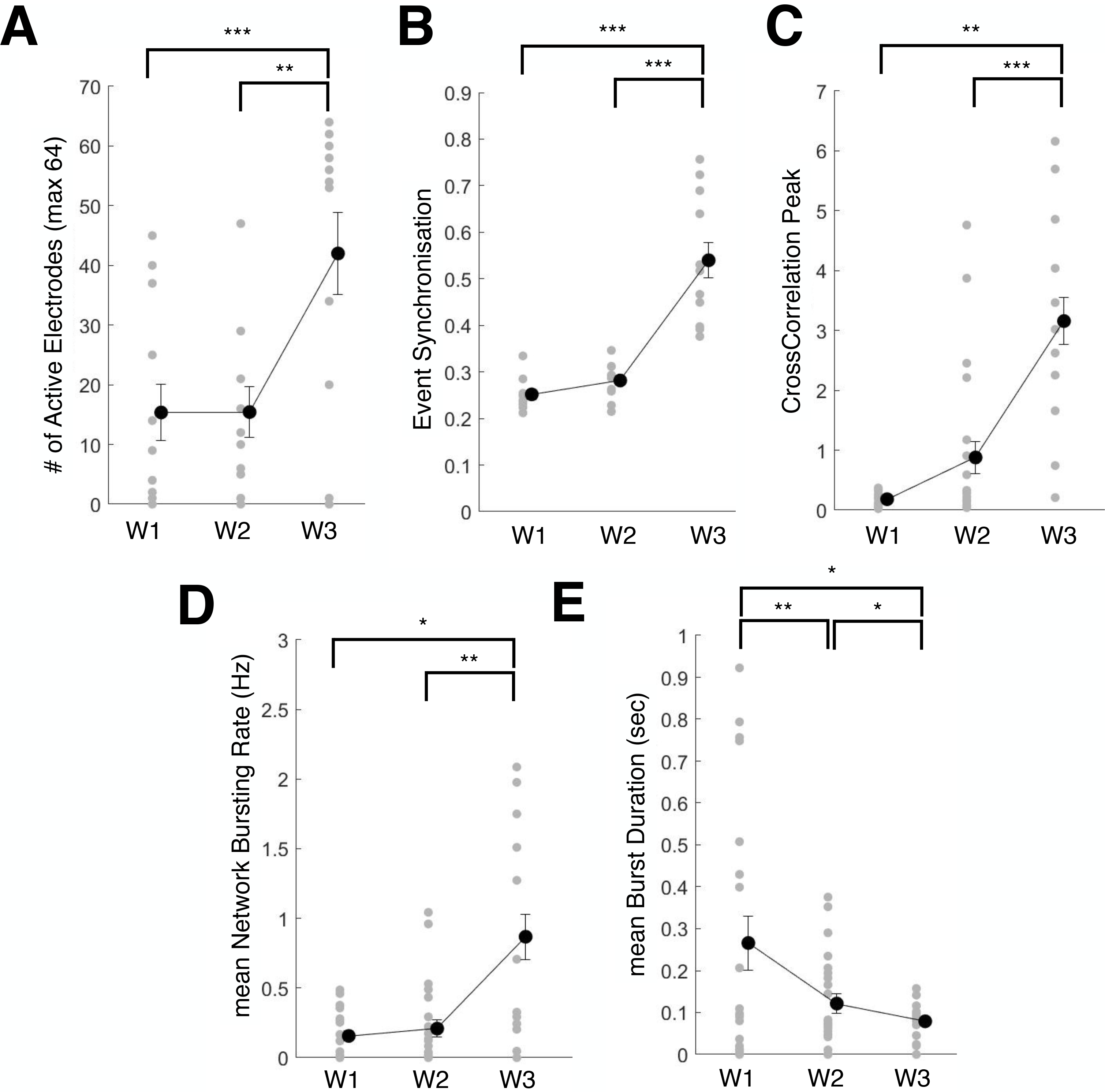
Temporal maturation of ESC network connectivity. (**A**) Change in number of active electrodes quantified from week 1 through week 3 in control condition. Data is shown scatter plot of individual well quantification and error bar plot using mean and s.e.m. * is for p<0.05, ** is for p<0.01 and *** is for p<0.001 using post-hoc multiple comparison with Holm-Sidak correction. (**B**) Change in Event Synchronization values from week 1 through week 3 in control condition. Data is shown as scatter plot of individual well quantification and error bar plot using mean and s.e.m. * is for p<0.05, ** is for p<0.01 and *** is for p<0.001 using post-hoc multiple comparison with Holm-Sidak correction. (**C**) Change in Cross-correlation Peak values from week 1 through week 3 in control condition. Data is shown as scatter plot of individual well quantification and error bar plot using mean and s.e.m. * is for p<0.05, ** is for p<0.01 and *** is for p<0.001 using post-hoc multiple comparison with Holm-Sidak correction. (**D**) Change in mean Network Bursting Rate values from week 1 through week 3 in control condition. Data is shown as scatter plot of individual well quantification and error bar plot using mean and s.e.m. * is for p<0.05, ** is for p<0.01 and *** is for p<0.001 using post-hoc multiple comparison with Holm-Sidak correction. (**E**) Change in mean Burst Duration values from week 1 through week 3 in control condition. Data is shown as scatter plot of individual well quantification and error bar plot using mean and s.e.m. * is for p<0.05, ** is for p<0.01 and *** is for p<0.001 using post-hoc multiple comparison with Holm-Sidak correction.

### L-glutamate treatment impaired network maturation excitability and synchrony

We then investigated the effect of L-glutamate exposure on the maturation and connectivity with neural networks derived from ESC-derived neurons and glia. L-glutamate treatment was administered 2 weeks post-seeding (Figure 4A; day14: 20 min exposure then wash; 100 μM). This dose was chosen to affect network maturation and plasticity without inducing complete silencing or critical impairment in network formation ^71^.

The immunohistochemical characterization of neuron/glial co-cultures following exposure to L-glutamate did not show any qualitative differences in comparison to control untreated cells. Both week 2 and week 3 immunohistochemical staining following L-glutamate exposure showed the presence of βIII-tub+ and SOX-1 + cells (Figure 4C). Similarly, O4+ and GFAP+ glial cells were also prominently evident within the co-cultures following L-glutamate treatment (Supplementary Figure 2B).

The individual characterization of each unit/electrode revealed a progressive increase in the firing rate (mFR: mean firing rate) and bursting rate (mBR: mean bursting rate) from week 2 to week 3 (Figure 4B). We quantified the rate of change from week 1 (baseline prior to L-glutamate treatment) for measures of excitability (# of bursting cells and wMFR) as well as measures of synchrony (Event synchronization and Cross-correlation Peak) by computing the z-scores at week 2 and week 3. We found that L-glutamate treated wells showed a significantly lower z-score for measures of excitability than the control wells (repeated-measure ANOVA, group: p < 0.05) particularly at week 2 (wMFR, p < 0.05, Student t-test) and at week 3 (# of cell bursting, p<0.05, Student t-test). The control wells had a proportion of wells that demonstrated a significant increase in excitability from week 1 (Z-score >1.96 or p <0.05) above 33% (range 33.3 to 64.3%) when compared to L-glutamate treated cells, which demonstrated lower excitability (<38%; range 0 to 37.5%) as indicated by the number of bursting cells and the wMFR (Figure 4D; well ratio).

Similarly, we found that L-glutamate treated wells showed a significantly reduced rate of change in synchrony measures (repeated-measure ANOVA, group: p < 0.05) particularly at week 2 (Event Synchronization, p < 0.05; Student t-test) and at week 3 (Event Synchronization, p < 0.05; Cross-correlation peak: p<0.01; Student t-test) when compared to control untreated cells. No significant differences were observed in the rate of network bursting (mBFR; repeated-measure ANOVA, group: p = 0.927). At week 3, we observed that L-glutamate treated cells demonstrated significantly lower measures of synchrony (Event Synchrony: 1/8 or 12.5%; Cross-correlation Peak: 4/8 or 50%) when compared to the untreated controls (Event Synchrony: 11/11 or 100%; Cross-correlation Peak: 10/11 or 90.1%).

Importantly, no differences were observed in the rate of change in the number of active electrodes between control and L-glutamate treated wells (repeated measure ANOVA, group x time: p = 0.298).

Altogether these results indicate that L-glutamate treated wells showed a reduced maturation in excitability and network connectivity immediately (immediately following L-glutamate treatment) and over a week following exposure, without affecting the rate of change in the number of active electrodes.

**Figure 4:**
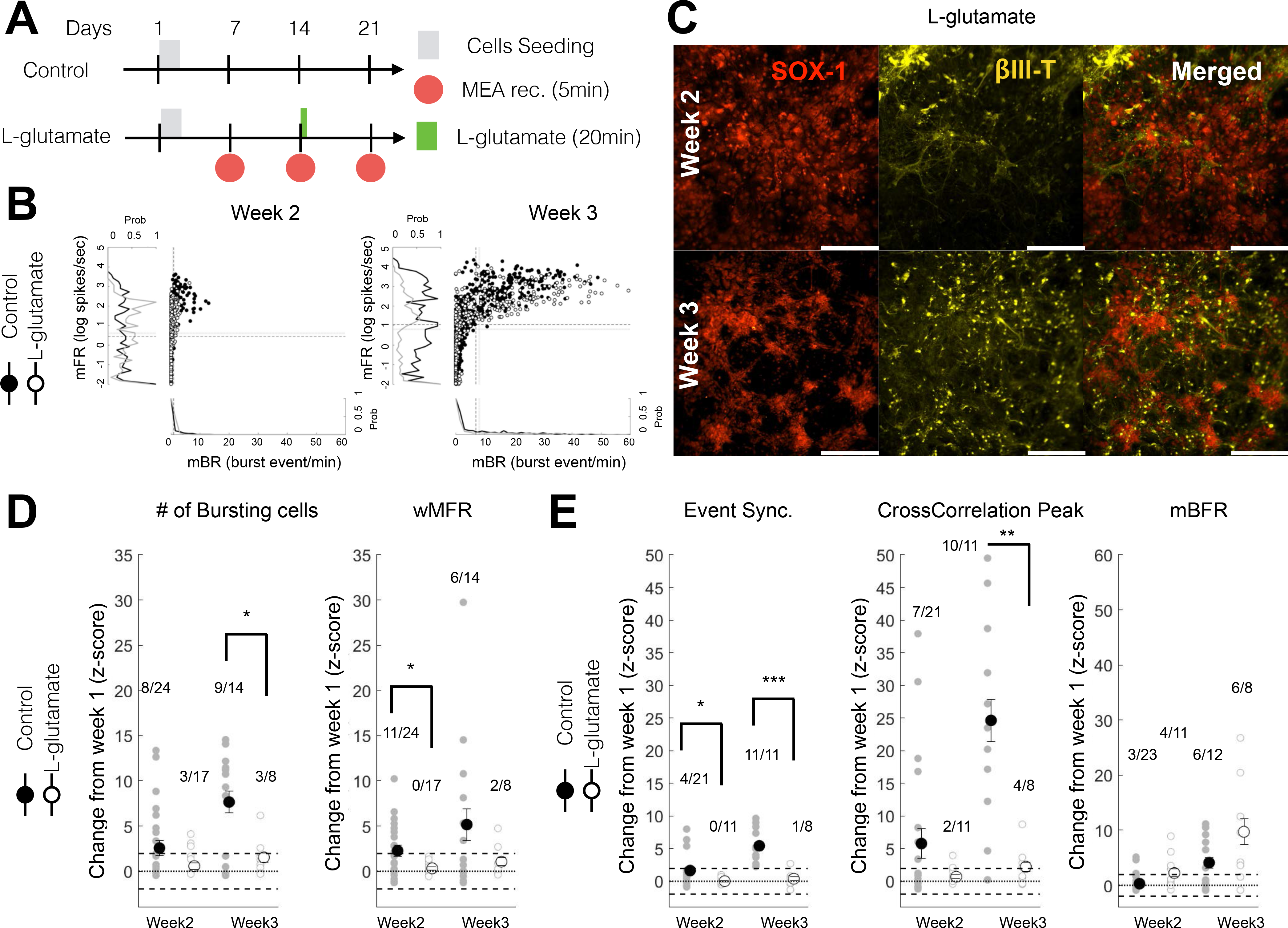
Acute L-glutamate treatment impaired network activity and synchrony immediately and over a week. (**A**) Experimental schedule for ESC-derived neuron seeding, culture and L-glutamate treatment. In the treatment group, 100μM of L-glutamate was given at day14 for 20min then washed out. Mea recording were performed from week 1 to week 3. (**B**) Scatter distribution of the mean burst firing rate (mBR; Burst event/min; x-axis) against the mean firing rate (mFR; log of spike/sec; y-axis) for individual unit recorded for the CTR (black filled circles) and L-glut (black open circle) groups. Data is shown for week 2 (left panel) and week 3 (right panel). The normalized distribution CTR (black line) and L-glut (gray line) groups are displayed as a projection of each axis. The dashed black line represents the average for the CTR group. The dotted black line represents the average for the L-glut group. mFR: mean firing rate for individual electrodes; mBR: mean bursting rate for individual electrodes. (**C**) ESC-derived neurons population for L-glutamate treatment (L-glut; 100 μM for 20 min at day 14) at week 2 (top panels) and week 3 (bottom panels). From left to right panels are shown SOX-1, pIII-Tubulin and merged image for all 2 fluorescent markers. The scale bar is for 200um. (**D**) Change at week 2 and week 3 (expressed as a Z-score from week1 baseline) of the number of bursting cells (left panel) and weighted mean firing rate (wMFR; right panel). For each group, the number of well with significantly increasing change over the total number of wells recorded is shown above the scatter plot. For post-hoc two-sample test *, ** and *** indicate p<0.05, p < 0.01 and p <0.001, respectively. wMFR: weighted mean population firing rate. (**E**) Change in network synchrony at week 2 and week 3 (expressed as a Z-score from week1 baseline) for event synchronization (left panel), cross-correlation peak (middle panel) and mean network burst firing rate (mBFR; right panel). For each group, the number of well with significantly increasing change over the total number of wells recorded is shown above the scatter plot. For post-hoc two-sample test *, ** and *** indicate p<0.05, p < 0.01 and p <0.001, respectively. mBFR: mean population burst firing rate.

### Electrical stimulation helps sustain network activity after exposure to L-glutamate

Since direct current stimulation (DCS) and low frequency stimulation (LFS) approaches have been demonstrated to induce functional recovery following CNS injury ^12,72–74^, we investigated the effects of electrical stimulation on the rate of change in the network excitability and connectivity after network inhibition by L-glutamate (Figure 4A and B).

First, we verified the direct effect of various electric stimulation protocols on the network responses (evoked response; Figure 5) on day 15 (where all stimulations protocol are delivered for the first time). For the “NoStim” condition, with or without L-glutamate infusion, an overall decrease in network activity was observed (Repeated measure ANOVA, time: p < 0.05; Figure 5A), which was particularly pronounced 15 min after the start of recording. LFS stimulation increased the network activity of 50% (1min binning; baseline 5 min pre-stimulation), up to the 15 minute mark. Post-LFS, a trend of increased activity was observed in comparison to NoStim condition, although these differences were not statistically significant. DCS stimulation showed evoked responses after L-glutamate infusion, during Stimulation and Post-stimulation, whereas Control condition did not show any effect. Combination DCS/LFS stimulation protocols with control and L-glutamate, similarly to LFS, induced a sustained network activity during stimulation and during stimulation/post-stimulation, respectively. A rhythmic response to LFS was observed in most wells (Figure 5B; 3 out of 4 wells; 100msec binning for wMRF). Representative peri-stimulus histograms illustrate the transient depression in population firing (wMFR z-score) followed by rebound of network activity stimulated by LFS protocols (Figure 5C).

**Figure 5:**
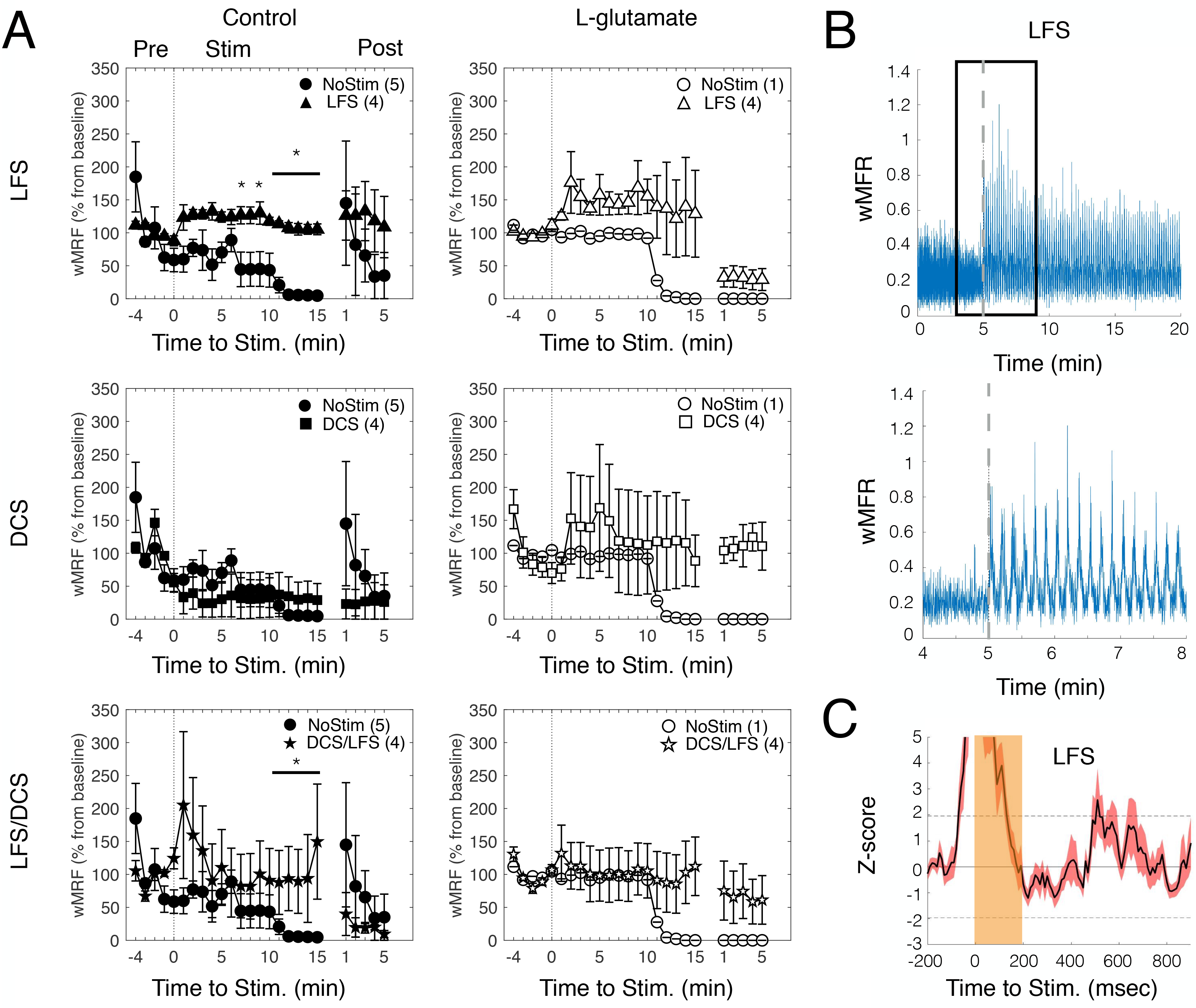
Evoked Network Response from different FES protocols. (**A**) Average wMFR as a percentage of the baseline (Pre) during the prestimulation (Pre, marked by a colored bar), stimulation (Stim) and poststimulation (Post) period. The dashed dark lines indicates the start of the 15min electrical stimulation. * indicate p< 0.05, ranksum between NoStim and Stimulation. (**B**) Representative trace showing the instantaneous wMFR (100 msec binning, stimulation duration 15min) for the LFS condition (upper panel; pulse of 200 msec duration at 0.1Hz, blue marks/bar) and a magnified representation of a 4min activity (lower panel; marked with a black box in the upper panel); dashed gray line indicate the start of stimulation. (**C**) Average peri-stimulus histogram of the population activity (pulse duration: 200 msec) expressed as a z-score (baseline: 200 msec prior stimulation; Average of 90 pulses). Note: rebound activity following activity suppression by the stimulation.

### Electrical stimulation initiates network recovery following L-glutamate induced inhibition

We then quantified changes at week-3 following the completion of all protocols (Figure 6A) relatively to week 1 (Z-score) in order to understand Long-term effect of FES on network maturation.

Following L-glutamate treatment, we found that LFS stimulation significantly increased the rate of maturation in neuronal excitability measures (# of cell bursting; one-way ANOVA, p = 0.039; LFS-LFS/DCS: p = 0.046; post-hoc Dunn correction) and marginally significant for wMFR (one-way ANOVA, p = 0.065; Sham-LFS: = 0.046; post-hoc Dunn correction) compared to the NoStim L-glutamate group (Figure 6C). Importantly, following L-glutamate, we found that LFS (Cell bursting and wMFR: 4/4 or 100%) and DCS (Cell bursting and wMFR: 3/5 or 60%) increased network excitability (Cell bursting: 4/8 or 50%; wMFR: 2/8 or 25%) when compared to NoStim. We observed that LFS and DCS after L-glutamate increased the rate of maturation in excitability within or higher than the range of NoStim Control, indicating a possible recovery to normal control level.

The DCS group showed an increasing trend with respect to event synchronization (one-way ANOVA, p = 0.364) and cross-correlation peak (one-way ANOVA, p = 0.665) measures of synchrony when compared to the NoStim group (Figure 6D), although, no significant group differences were observed. The DCS group also showed a marginally significant difference in mBFR when compared to the NoStim group (one-way ANOVA, p = 0.053; Figure 6D, right panel), with a similar trend to the other measures of synchrony as described above. As for measure of excitability, LFS (Cell bursting: 3/4 or 75%; wMFR: 4/4 or 100%) and DCS (Cell bursting: 5/5 or 100%; wMFR: 3/5 or 60%) groups induced greater cell bursting and wMFR when compared to the NoStim group (Cell bursting: 5/8 or 62.5%; wMFR: 4/8 or 50%). Combination LFS/DCS treatment did not demonstrate any clear effects on these measures.

Altogether, these results indicate that LFS and DCS had a positive impact on the rate of change in excitability and synchrony compared to the NoStim L-glutamate groups and returning within range or higher than NoStim control levels.

**Figure 6:**
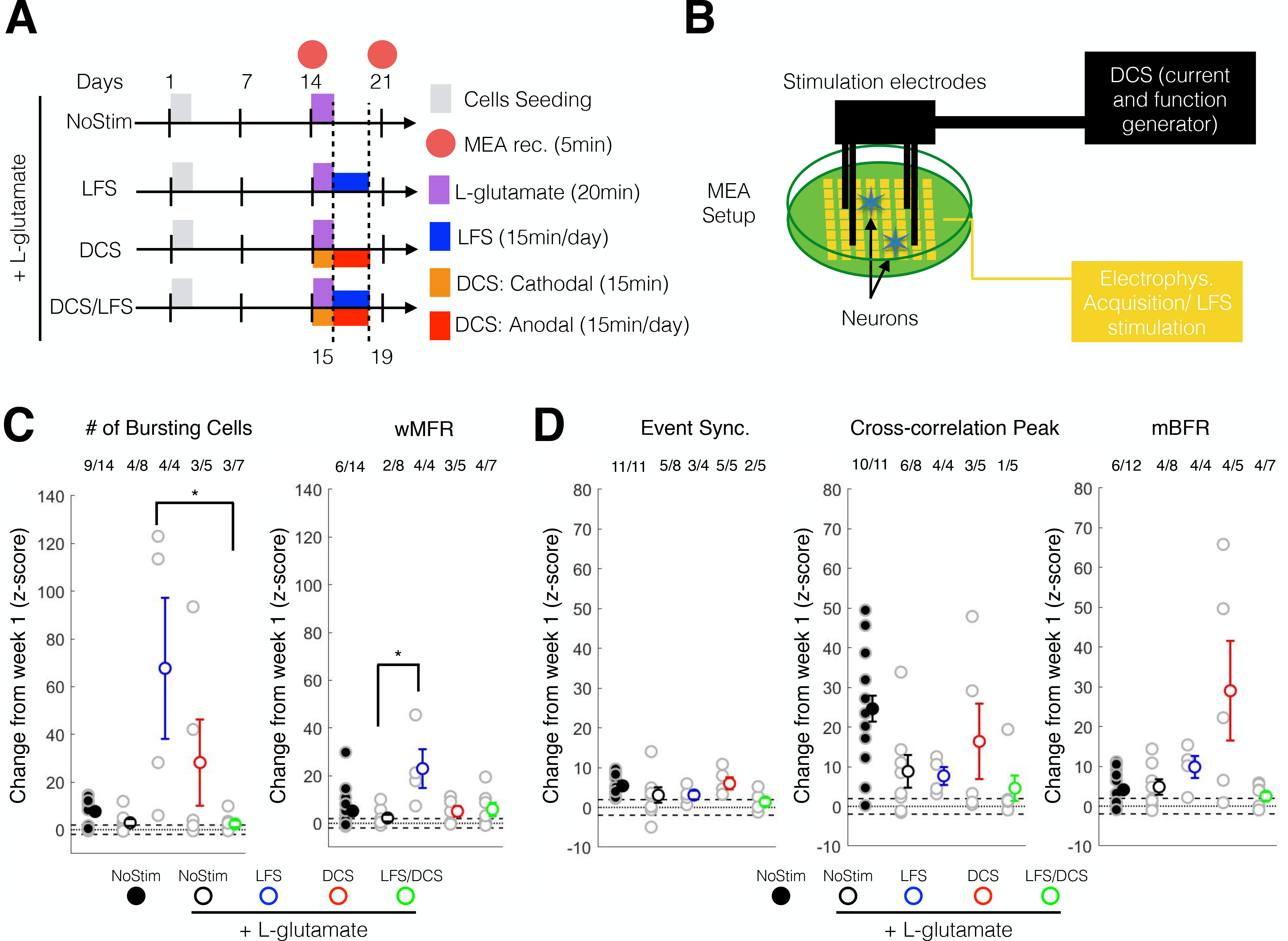
DCS/LFS stimulation enhanced the maturation in excitability and neural synchrony following acute L-glutamate treatment. (**A**) Experimental schedule for ESC-derived neuron seeding, culture and treatment. L-glutamate (100μM) was given at day14 for 20min then washed out. Low frequency stimulation (LFS) was given from day 15 to day 19 using 10 μA at 0.1Hz, 15min/day. Direct current stimulations were given in two phases: day14 a cathodal stimulation was performed for 15min; from day 15 to day 19, anodal stimulation were given for 15 min per day. Recording was performed on day 14 (week 2) and day 21 (week 3). (**B**) Schematic set up for electrical stimulation on MEA plate. LFS stimulation (10μA, 0.1Hz, 15min/day) where delivered through the MEA electrode using theMaestro systems. Four stainless screws were positioned above the MEA cultured neurons and delivered a controlled current (DCS: single-time 10μA monophonic cathodal 15min and daily 10μA monophonic anodal current, 15min/day) using a custom battery-powered system. (**C**) Change at week 3 (expressed as a Z-score from week1 baseline) of the number of bursting cells (left panel) and wMFR (right panel). For post-hoc multiple comparison using Dunn-Sidak correction *, ** and *** indicate p<0.05, p < 0.01 and p <0.001, respectively. wMFR: weighted mean population firing rate. (**D**) Change at week 3 (expressed as a Z-score from week1 baseline) of the event synchronization (left panel), cross-correlation peak (middle panel) and mBFR (right panel). For post-hoc multiple comparison using Dunn-Sidak correction *, ** and *** indicate p<0.05, p < 0.01 and p <0.001, respectively. mBFR: mean population burst firing rate.

### Electrical stimulation following L-glutamate-inhibition significantly enhanced excitability and plasticity related gene expression

In order to understand the underlying effects of electrical stimulation of untreated cells and following L-glutamate treatment, we investigated the change in expression of transcripts encoding N-methyl-D-aspartate (NMDA) receptor sub unit A and B (*NR2A* and *NR2B*; transcripts encoding two receptors linked to glutamate excitation and the modulation of long-term potentiation and depression), brain-derived neurotrophic factor (*BDNF*; transcript encoding a protein that promotes synaptogenesis and neurogenesis) and RAS-related protein (*RAB3A*; transcript encoding a protein related to synaptic vesicles transport).

The quantitative real-time PCR analysis (fold up and down) showed that under the control condition (no L-glutamate treatment), LFS and DCS stimulation induced a reduction in *NR2A* (LFS: p = 0.270; DCS: p = 0.002) and *NR2B* (LFS: p < 0.001; DCS: p < 0.001) expression when compared to the unstimulated controls (Control NoStim). *RAB3A* expression was significantly increased under LFS (p < 0.001) and decreased for the combination of LFS/DCS stimulation (p = 0.001) when compared to the Control NoStim (Figure 7A). In contrast to LFS and DCS stimulated groups, the cells exposed to combination LFS/DCS stimulation induced significantly increased expression of *NR2B* encoding transcripts (p = 0.002) when compared to the Control NoStim.

Following L-glutamate treatment, we found that all transcripts encoding excitability and plasticity related proteins were significantly downregulated in the L-glutamate NoStim group when compared to the Control NoStim group (p < 0.01; Figure 7B), with an ~8 fold downregulation of *NR2A* (RQ: −8.6±0.78; p < 0.001). In contrast, L-glutamate treated cells subjected to electrical stimulation (LFS, DCS, and combination LFS/DCS) significantly upregulated *NR2A* expression (LFS p < 0.001; DCS p < 0.001; LFS/DCS p = 0.012) when compared to L-glutamate NoStim treated controls (Figure 7C). Interestingly, the gene expression of *NR2B* was significantly decreased in all cells subjected to stimulation protocols (LFS: p < 0.001; DCS: p = 0.457; LFS/DCS: p < 0.001) when compared to L-glutamate NoStim. LFS and DCS also significantly increased the expression of *BDNF* (LFS and DCS: p < 0.001) and *RAB3A* (LFS: p = 0.019; DCS: p < 0.001) following L-glutamate treatment when compared to L-glutamate NoStim, which was in stark contrast to combination LFS/DCS treated group that demonstrated a significant decrease in *BDNF* expression.

In order to better understand the effect of stimulation following L-glutamate treatment, we tested the relative difference in fold change of expression between each stimulation protocol with and without L-glutamate. For all genes except *NR2B*, we detected a significant effect for main factor treatment (two-way ANOVA, treatment: p < 0.05; two levels: control and L-glutamate) and main factor interaction (two-way ANOVA, treatment x stimulation: p < 0.005: stimulation, 3-levels: LFS, DCS and LFS/DCS). *NR2B* only had a significant factor interaction (two-way ANOVA, treatment x stimulation: p < 0.001) and a marginal effect of the main factor treatment (two-way ANOVA, treatment: p = 0.054). These results are consistent with the L-glutamate induced significant downregulation (Figure 7B), and the subsequent stimulation-induced significant upregulation of these transcripts (Figure 7C). Post-hoc comparison between control and L-glutamate condition revealed a significant increase for *NR2A* for all stimulation conditions (p< 0.001, Holm-Sidak correction). *NR2B* expression was significantly decreased under L-glutamate treatment compared to control for DCS (P < 0.05, Holm-Sidak correction) and DCS/LFS (P < 0.001, Holm-Sidak correction) groups. *BDNF* expression was significantly increased under L-glutamate treatment compared to control for DCS and DCS/LFS (P < 0.001, Holm-Sidak correction) groups. Under L-glutamate treatment, only the DCS condition evoked the increased expression of *RAB3A* when compared to control.

Notably, following L-glutamate (RQ to L-glutamate NoStim), post-hoc multiple comparison indicated that LFS and DCS induced significantly higher expression of *NR2A* (p < 0.05, Holm-Sidak correction), *BDNF* (p < 0.001, Holm-Sidak correction) and *RAB3A* (p < 0.001, Holm-Sidak correction) than the combination LFS/DCS group. The expression of *NR2B*, however, was significantly decreased by the combination LFS/DCS (p < 0.05, Holm-Sidak correction) following L-glutamate exposure.

Altogether these results indicate that the DCS and LFS alone following L-glutamate treatment induced the significant up-regulation of excitability (*NR2A*) and plasticity (*BDNF* and *RAB3A*) related genes, whereas LFS/DCS combination had adverse effects.

**Figure 5:**
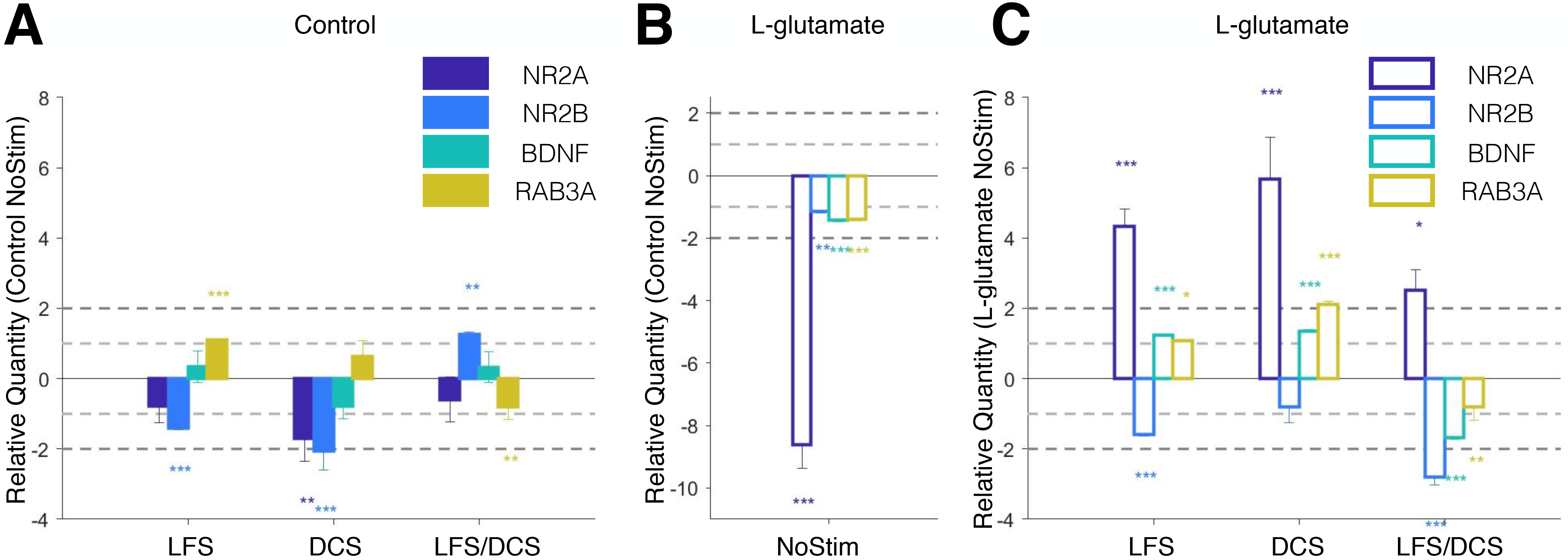
DCS/LFS stimulation alters the expression profile in NR2A, NR2B, BDNF and RAB3. (**A**) Relative quantity (RQ to Control NoStim) for gene expression change for LFS, DCS and LFS/DCS stimulation without toxicity at week 3. Data is shown as mean +/− SEM. *, ** and *** are for p<0.05, p<0.01 and p<0.001, respectively, on two-sample student t-test between test and control samples (n = 6 repeats per group). Light and dark gray dashed lines mark 1- and 2-fold change in expression. NR2A: NMDA receptor sub unit 2A, NR2B: NMDA receptor sub unit 2B, BDNF: brain-derived neurotrophic factor, RAB3A: RAS-related protein rab3. (**B**) Relative quantity (RQ to Control NoStim) for gene expression change following L-glutamate toxicity at week 3 (L-glutamate NoStim). Data is shown as mean +/− SEM. *, ** and *** are for p<0.05, p<0.01 and p<0.001, respectively, on two-sample student t-test between test and control samples (n = 6 repeats per group). Light and dark gray dashed lines mark 1- and 2-fold change in expression. NR2A: NMDA receptor sub unit 2A, NR2B: NMDA receptor sub unit 2B, BDNF: brain-derived neurotrophic factor, RAB3A: RAS-related protein rab3. (**C**) Relative quantity (RQ to L-glutamate NoStim) for gene expression change for LFS, DCS and LFS/DCS stimulation following L-glutamate toxicity at week 3. Data is shown as mean +/− SEM. *, ** and *** are for p<0.05, p<0.01 and p<0.001, respectively, on two-sample student t-test between test and control samples (n = 6 repeats per group). Light and dark gray dashed lines mark 1- and 2 fold change in expression. NR2A: NMDA receptor sub unit 2A, NR2B: NMDA receptor sub unit 2B, BDNF: brain-derived neurotrophic factor, RAB3A: RAS-related protein rab3.

## DISCUSSION

In our current work, we present a first evaluation of direct current stimulation protocols on the maturation of neuronal networks following acute glutamate exposure. We demonstrated using ESC-derived neuron and glia co-cultures on MEAs, that the glutamate-induced impairment in maturation (Figure 4) can be overcome using direct current stimulation (DCS and LFS; Figure 6) through potential mechanisms that involve the change in expression of excitability and plasticity-related genes NR2A/B, BDNF and RAB3A (Figure 7). These studies provide: 1) A proof-of-concept of the utility of MEAs as a platform to investigate the effects of FES following excitotoxic agents in physiologically relevant conditions; and 2) validation that DCS and LFS can be used to evoke network maturation and recovery following acute glutamate treatment.

### Model for glutamate-induced excitotoxicity in ESC-derived co-cultures

Glutamate is one of the most common excitotoxic agents responsible for the propagation of secondary neuronal damage after TBI ^75^ Acute glutamate release (>300μMol) reportedly results in cell swelling and neurotoxicity due to NMDA-receptor binding ^76,77^ and the failure of glutamate re-uptake by astrocytes ^78,79^ in cortical neuron and glial co-cultures. Previous MEA-based studies showed a dose-dependency to L-glutamate in network response and mostly focused on a single time point change in network excitability or connectivity ^44,71,80–82^. Our results corroborate these previous findings, and further demonstrate that acute glutamate treatment (single dose 100 μM of L-glutamate for 20 min) resulted in reduced network rate of maturation in excitability (Figure 4C) and connectivity (Figure 4D) in ESC-derived neuron and glial co-cultures, similarly to embryonic cortical neuron and glial co-cultures, without inducing any significant difference in the change in number of active electrodes at week 3 (i.e. lesser impact on the viability of the neural cells; Supplementary Figure 4; repeated-measure ANOVA, group × time: p = 0.298) or silencing ^71^. Acute exposure to L-glutamate might cause delayed and hour-long increase in intracellular calcium concentration ^83^ and in turn might modulate plasticity changes ^84,85^. In this study, we have successfully demonstrated immediate (within minutes) and extended (1 week) changes in network rate of maturation following acute glutamate exposure (Figure 4) and its corresponding long-term modulation on plasticity-related gene expression (Figure 7B), which to our knowledge, has not been shown previously. These results validate the application of MEA recording/stimulation platforms for the high-throughput and longitudinal assessment of cellular and neuronal network maturation, and correlative genomic expression in response to excitotoxic stimuli *in vitro*. ^86,87^

### Acute L-glutamate and NR2A/B expression

NMDA subunit composition and postsynaptic location of NMDARs are critical determinants of synaptic plasticity ^88^ NR2A receptors are synaptically located and associated with long-term potentiation (LTP), while NR2B receptors are extrasynaptically located associated with long-term depression (LTD). Most importantly, the ratio of NR2A/NR2B is thought to be predictive of plasticity changes as observed in brain development studies ^89–91^. We observed that in our control condition, the general expression level of *NR2A* and *NR2B* transcripts decreased following LFS and DCS, whereas combination DCS/LFS increased *NR2B* expression. Importantly, following acute L-glutamate treatment, the L-glutamate treated NoStim group significantly down-regulated all transcripts, and particularly NR2A in comparison to the control NoStim condition (Figure 7B), which might be consistent with a NMDA receptor-mediated calcium influx and neuronal over-excitation ^92^ Interestingly, we found a significant up-regulating (*NR2A, BDNF* and *RABA*) and down-regulating (*NR2B*) effect of stimulation when coupled with L-glutamate treatment (Figure 7; two-way ANOVA, treatment factor: p < 0.001; interaction treatment × stimulation: p < 0.004). Indeed, both LFS and DCS resulted in a significant up-regulation of *NR2A* (p < 0.001) expression and down-regulation of *NR2B* (LFS: p < 0.001) compared to L-glutamate NoStim condition, which was consistent with the observed improvement in the network maturation over one week period (week3). These results support the use of LFS and DCS post injury-induced neurotoxicity for the recovery of neuronal network ^12,74,93^, but also provide evidence that the preventive use of these stimulation regimens could protect from neurotoxicity by down-regulating NMDA-receptor expression ^92,94,95^.

### DCS and LFS likely improved network maturation following glutamate-induced impairment by upregulating plasticity genes

Cathodal/anodal tDCS and LFS are two of the most common non-invasive stimulation approaches in the pre-clinical and clinical literature ^1^ for the therapeutic treatment of brain injury and neurodegenerative diseases. These two protocols have not been extensively compared and their mechanisms of action for treating brain disorders/injury are poorly understood. We found that LFS significantly increased the rate of maturation in excitability (# of cell bursting: p < 0.05, and wMFR: p = 0.065; 100% of well showing increase versus < 50% for the sham group) and also promoted a marginal increase in synchrony maturation as seen through cross-correlation peak and mBFR (mean bursting firing rate; p = 0.05; 100% of well significantly increasing versus <75% for the sham group). LFS has been shown to favor long-term potentiation (LTP) through the Trek/BDNF pathway ^16^. At the electrophysiological level, we found that LFS could help sustain network activity (Figure 5A) during stimulation, which effect did not persist post-stimulation (Figure 5A, Post-stimulation period). Interestingly, LFS alone in the control condition promoted *RAB3A* expression without inducing a corresponding increase in *BDNF* expression (Figure 7A). However, following L-glutamate treatment, LFS significantly up-regulated both *BDNF* and *RAB3A* (Figure 7B), which also corresponded to an improved network connectivity as seen through our synchrony measures (Figure 6D). *RAB3A* is an intracellular vesicular trafficking protein required for calcium exocytosis and is thought to be required (i.e. up-stream) for the BDNF-dependent increase in synaptic plasticity ^96^ Therefore, RAB3A-BDNF might corroborate the effect of LFS on the plastic change observed following L-glutamate excitotoxicity, although those changes under control conditions does not support previous observations ^16^

Similarly to LFS, we observed that DCS following L-glutamate treatment showed a trend of increased excitability (Figure 4C; # of bursting cells) and a clear, although non-significant, increase in rate of maturation in synchrony (Figure 4D; Cross-correlation peak and mBFR) as well as increased *BDNF* and *RAB3A* expression. Surprisingly, DCS in control condition induced the significantly reduced expression of *NR2A* and *NR2B* expression without a specific effect on *BDNF* or *RAB3A* (Figure 7A) and no significant effect on network maturation (oneway ANOVA, p > 0.05 for all measures; Supplementary Figure 4BC). The latter result might come in contrast with our observation that DCS might induce long-lasting activation of neural network (Figure 5A) and previous *in vivo* studies that suggest a possible increase in plasticity and memory through an epigenetic modulation of BDNF expression by tDCS ^65,97^, although it is important to note that the DCS stimulation used in this study was a combination of cathodal and chronic anodal stimulations and was tailored for a use following glutamate treatment as opposed to one-time anodal excitatory stimulation ^65^ Overall, our results confirm that the therapeutic use of LFS and DCS mechanistically contributes to the enhancement of network maturation consistently with an up-regulation of plasticity gene expression.

### DCS/LFS combination demonstrated negative effects on molecular and network plasticity following L-glutamate treatment

Since DCS and LFS are two promising neurostimulation methods for the modulation of learning and memory ^98^, we sought to test the combination of the two protocols on MEA neural cell co-cultures. Surprisingly, in control condition combining DCS and LFS resulted in down-regulation of *BDNF* and an up-regulation of *NR2B* expression supported by no significant change in network synchrony or excitability (one-way ANOVA, p > 0.05; Supplementary Figure 4C). Following acute L-glutamate, DCS/LFS combination increased *NR2A* and down-regulated *NR2B* expression, in addition to strongly suppressing *BDNF* and *RAB3A* expression. Consistent with our qRT-PCR results, following L-glutamate treatment, DCS/LFS did not show change in the rate of maturation in excitability (Figure 6D) although evoked responses from the network were observed during and poststimulation (Figure 5A). The individual LFS and DCS parameters used in these studies were informed from previous works ^16,98^. These parameters when used in combination DCS/LFS combination may have induced neuronal over-excitation (Figure 5A) and fatigue leading to the negative results observed. Future studies shall include a better correspondence and comparison of electrical stimulation methods ^46^, which would in turn likely provide optimal stimulation parameters for *in vivo* applications.

**Table 1.** Pre-validated mouse qRT-PCR primers used to measure gene expression at week 3.

| Gene | Symbol | Refseq Number |
| --- | --- | --- |
| *NR2A* | Grin2a | NM_008170 |
| *NR2B* | Grin2ab | NM_008171 |
| *BDNF* | BDNF | NM_001048139 |
| *RAB3A* | RAB3A | NM_001166399 |
| *GAPDH* | GAPDH | NM_008084 |
| *HPRT1* | HPRT | NM_013556 |

## FUNDING AND ACKNOWLEDGEMENTS

This research was supported by startup funds from UGA, seed grant funding from the Georgia Institute of Technology/Emory University Regenerative Engineering and Medicine Center, NIH R01(1R01NS099596-01A1) award to Lohitash Karumbaiah and National Science Foundation Science and Technology Center for Emergent Behaviors of Integrated Cellular Systems, Grant CBET-0939511 to Steve Stice.

## Author Contribution

C.-F. L. performed stimulation assays of ESC-derived neurons with and without acute exposure to L-glutamate, analyzed the data and prepared all the figures. L.J. performed all MEA experiments and immunohistochemistry. L.K. performed all qRT-PCR experiments. M.S. designed the DC stimulation apparatus. C.-F. L. and L.K. wrote the manuscript. K.T., L.M., S.S., M.G. and L.K. edited the manuscript.

## Supplementary Figures

**Supplementary Figure 1:**
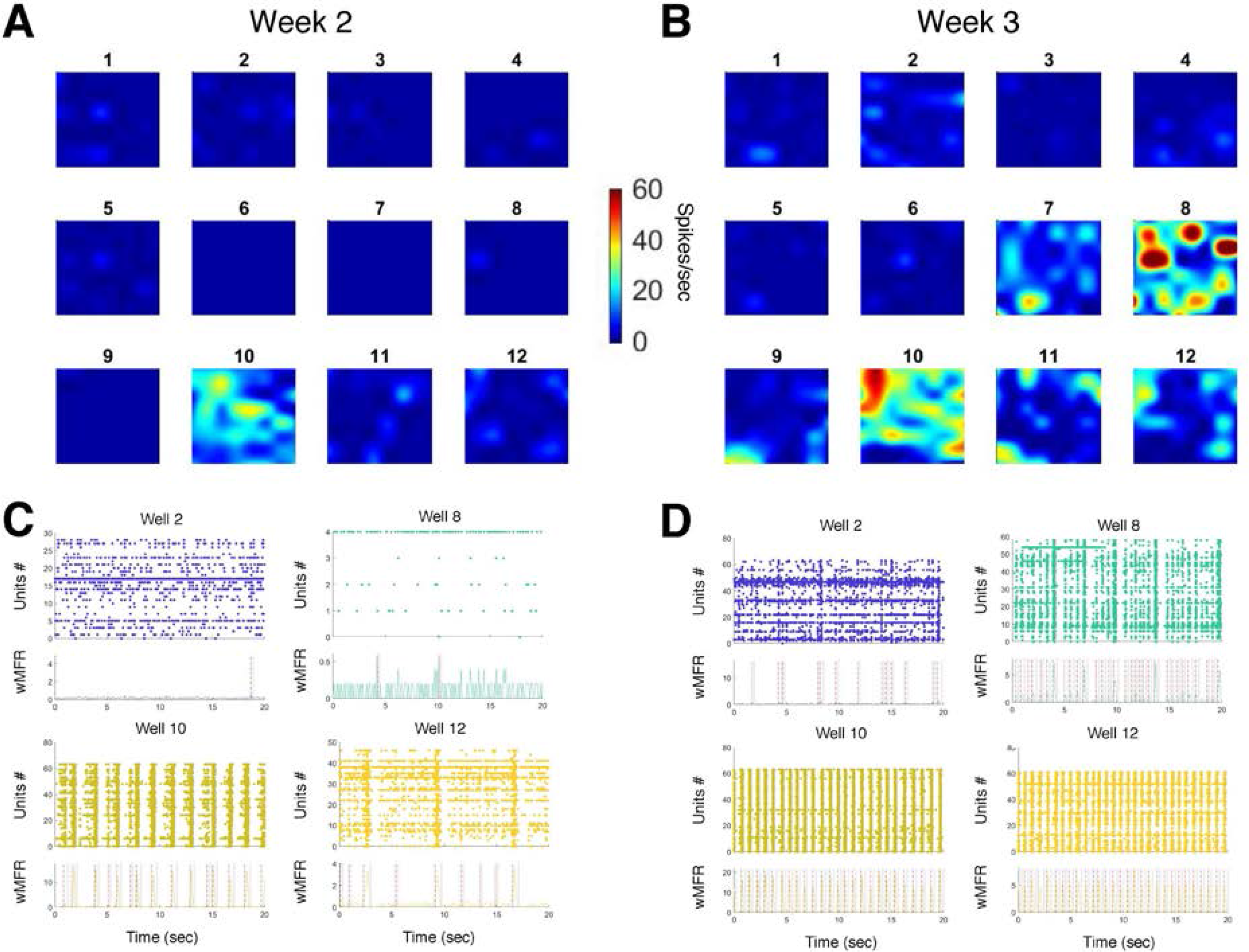
Representative network activity in control condition. (**A**) Average activity map (spikes/sec) for 12 wells recorded under control conditions on week 2. Average activity is estimated for each electrode separately over a period of 5min. (**B**) Average activity map (spikes/sec) for 12 wells recorded under control conditions on week 3. Average activity is estimated for each electrode separately over a period of 5min. (**C**) Raster plot of 4 wells activity recorded at week 2 (top panel) and the wMFR (bottom panel; spikes/bin/unit; bin size: 100 msec); 20-sec epoch. The vertical dashed and dotted lines indicate the start and stop of a detected population burst, respectively. (**D**) Raster plot of 4 wells activity recorded at week 3 (top panel) and the wMFR (bottom panel; spikes/bin/unit; bin size: 100 msec); 20-sec epoch. The vertical dashed and dotted lines indicate the start and stop of a detected population burst, respectively.

**Supplementary Figure 2:**
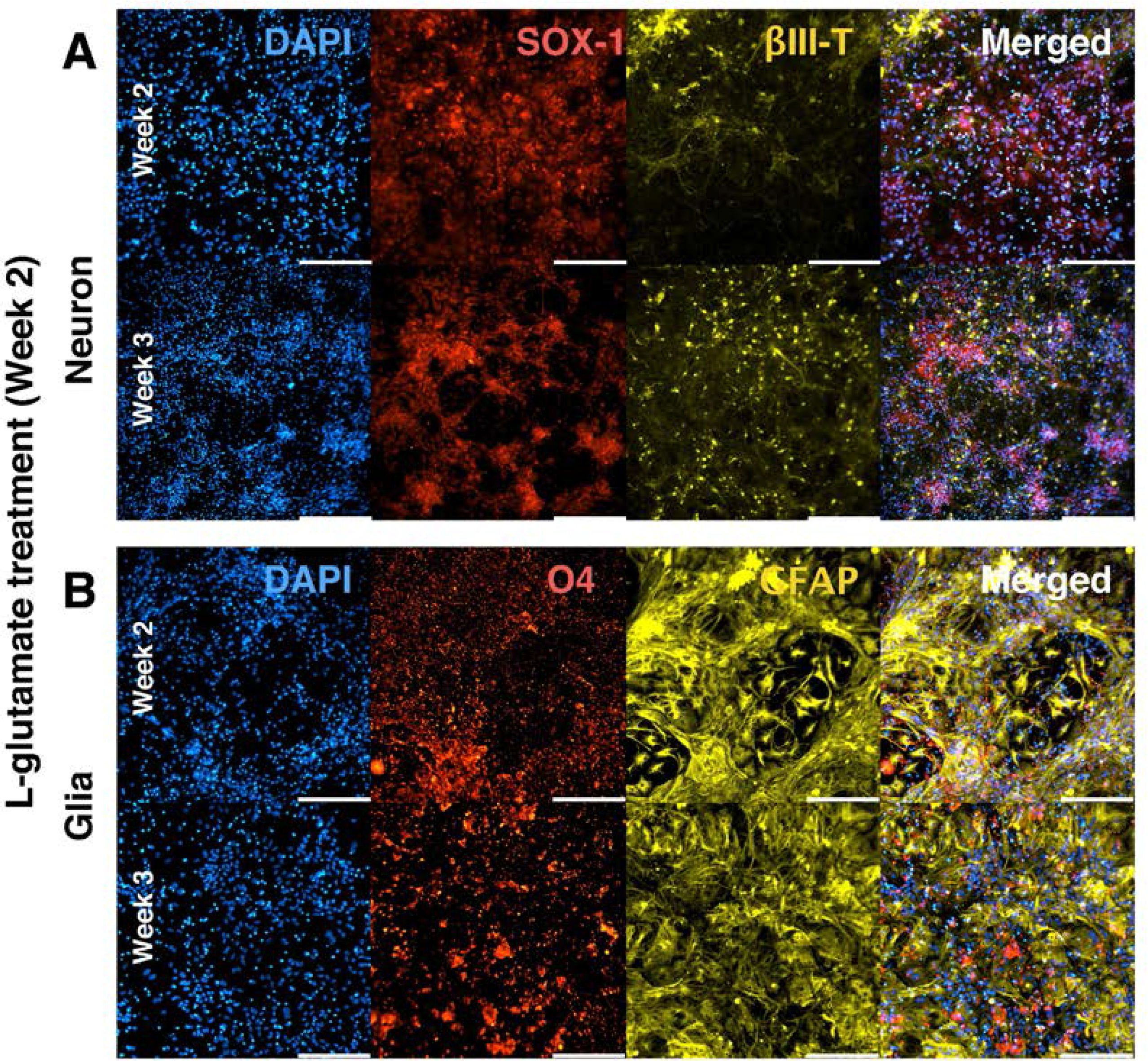
ESC-derived neurons and glia matured into neural tissue after L-glutamate cytotoxicity (week2) (**A**) ESC-derived neurons population for the L-glutamate treated group at week 2 (top panels; immediately after cytotoxicity delivery) and week 3 (bottom panels). From left to right panels are shown DAPI, SOX-1 (Neural progenitor origin marker), βIII-Tubulin (Neuronal marker) and merged image for all 3 fluorescent markers. The scale bar is for 200 μm. (**B**) ESC-derived glial population for the L-glutamate treated group at week 2 (top panels; immediately after cytotoxicity delivery) and week 3(bottom panels). From left to right panels are shown DAPI, O4 (Oligodendrocyte marker), GFAP (Glial marker) and merged image for all 3 fluorescent markers. The scale bar is for 200 μm.

**Supplementary Figure 3:**
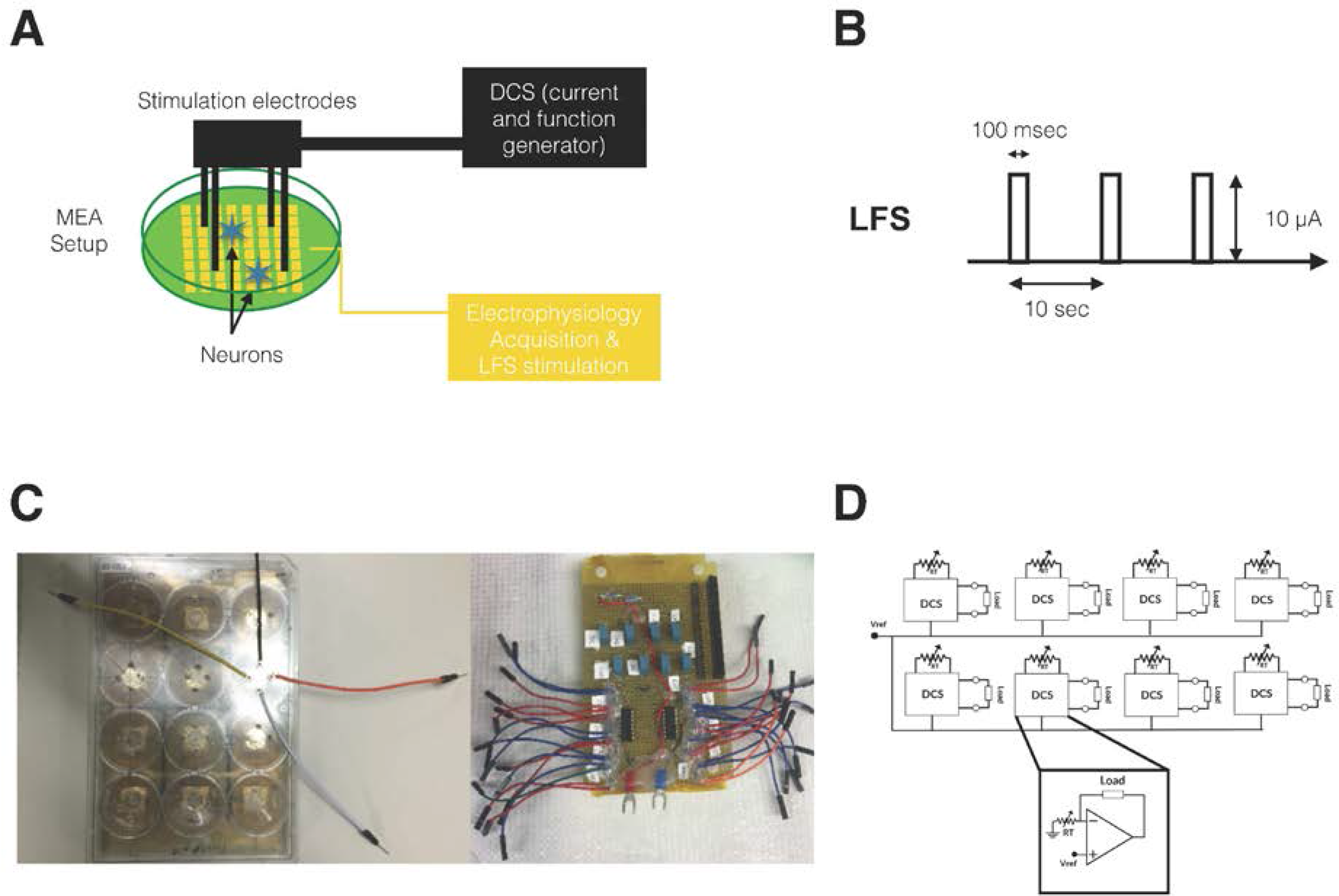
*In vitro* stimulation set up. (**A**) Schematic setup for electrical stimulation on MEA plate. LFS stimulation (10 μA, 0.1 Hz, 15 min/day) where delivered through the MEA electrode using the Maestro systems. Four stainless steel screws were positioned above the MEA cultured neurons and delivered a controlled current (DCS: 10 μA monophasic cathodal 15 min and 10 uA monophasic anodal current, 15 min/day) using a custom battery-powered system. (**B**) Parameters used for the LFS protocol. LFS was delivered at 0.1 Hz at an intensity of 10uA for 15 min every day for 5 days. The pulse width was set to 100 msec. (**C**) Custom-made DCS delivery system. The MEA cover plate was customized to host 4 stainless screws for DCS delivery (left panel) with a current distribution controlled through a microcontroller (right panel). (**D**) Block diagram of the custom eight-channel direct current stimulation (DCS) system. The design supports the control of current passing through the load (i.e. a mixed population of neurons and glia) using a variable resistor (RT). The schematic of DCS block is shown in inset which utilizes a negative feedback on the inverting input of the op-amp (MC33204P). As a result, the amount of current passing through the load is proportional to V_ref_. Using this configuration, anodal and cathodal currents were set by changing the polarity of load.

**Supplementary Figure 4:**
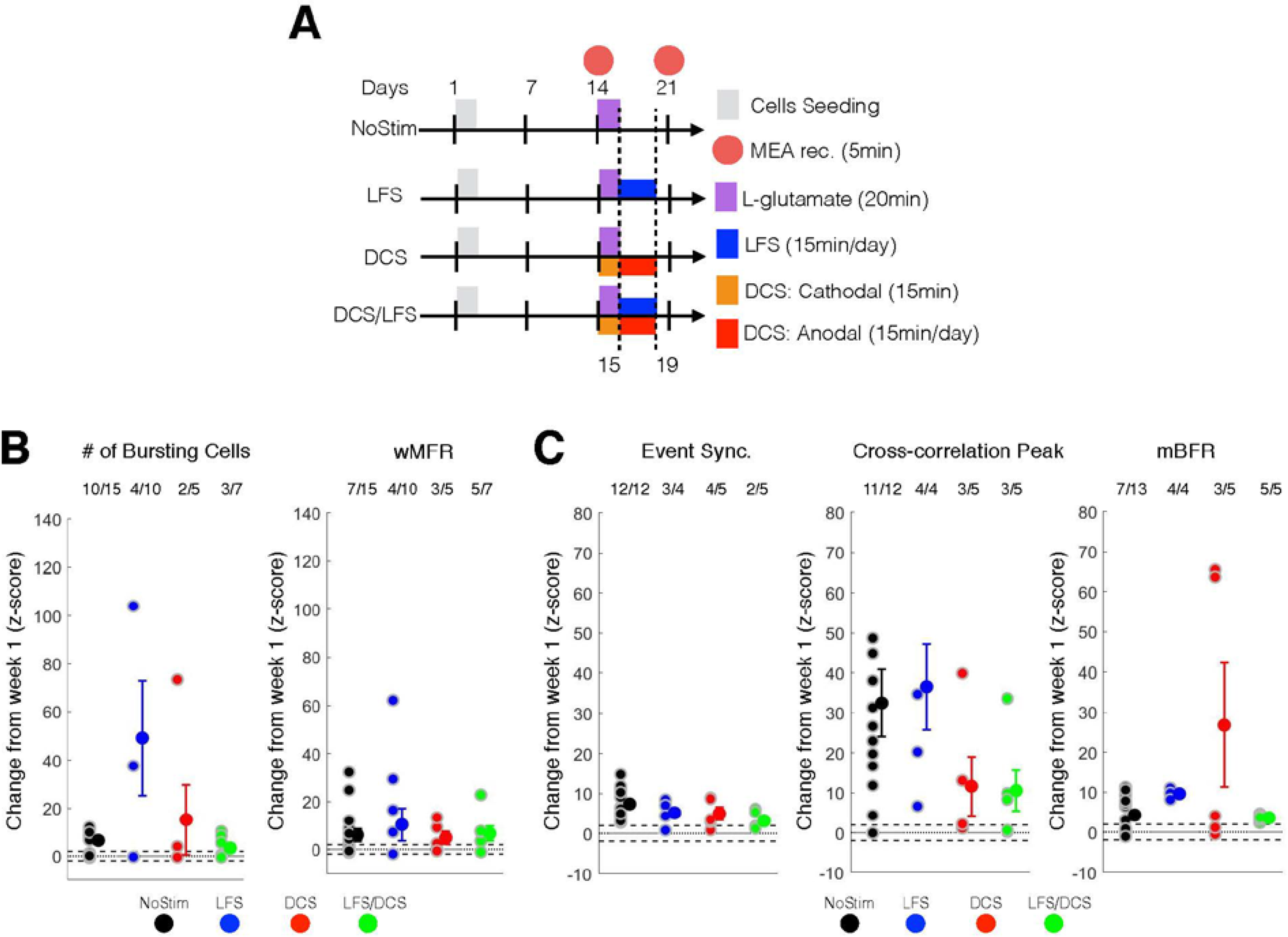
DCS/LFS stimulation did not affect maturation in control condition. (**A**) Experimental schedule for ESC-derived neuron seeding, culture and treatment. (**B**) Change at week 3 (expressed as a Z-score from week1 baseline) of the number of bursting cells (left panel) and wMFR (right panel) for the control condition. For post-hoc multiple comparison using Dunn-Sidak correction, ** and *** indicate p<0.05, p < 0.01 and p <0.001, respectively. wMFR: weighted mean population firing rate. (**C**) Change at week 3 (expressed as a Z-score from week1 baseline) of the event synchronization (left panel), cross-correlation peak (middle panel) and mBFR (right panel) for the control condition. For post-hoc multiple comparison using Dunn-Sidak correction *, ** and *** indicate p<0.05, p < 0.01 and p <0.001, respectively. mBFR: mean population burst firing rate.

